# Voxel Carving Based 3D Reconstruction of Sorghum Identifies Genetic Determinants of Radiation Interception Efficiency

**DOI:** 10.1101/2020.04.06.028605

**Authors:** Mathieu Gaillard, Chenyong Miao, James C. Schnable, Bedrich Benes

## Abstract

Changes in canopy architecture traits have been shown to contribute to yield increases. Optimizing both light interception and radiation use efficiency of agricultural crop canopies will be essential to meeting growing needs for food. Canopy architecture is inherently 3D, but many approaches to measuring canopy architecture component traits treat the canopy as a two dimensional structure in order to make large scale measurement, selective breeding, and gene identification logistically feasible. We develop a high throughput voxel carving strategy to reconstruct three dimensional representations of maize and sorghum from a small number of RGB photos. This approach was employed to generate three dimensional reconstructions of a sorghum association population at the late vegetative stage of development. Light interception parameters estimated from these reconstructions enabled the identification of both known and previously unreported loci controlling light interception efficiency in sorghum. The approach described here is generalizable and scalable and it enables 3D reconstructions from existing plant high throughput phenotyping datasets. For future datasets we propose a set of best practices to increase the accuracy of three dimensional reconstructions.

## 1 INTRODUCTION

Yields of agricultural plants will need to grow 70% by 2050 in order to meet growing demands for food and feed, which are projected to double between 2005 and 2050 (Tilman et al., 2011; Alexandratos and Bruinsma, 2012). Currently we are not on track to meet this demand. In many parts of the world increases in wheat and rice yields have dropped to zero as observed by Grassini et al. (2013). While maize yields continue to increase, the increased annual spending on breeding is required each year in order to achieve the same fixed annual increase in yields per year (Grassini et al., 2013). Developing new high yielding and more resilient crop varieties depends on the ability to score large populations of new lines for traits. All else constant, the more new data can be collected from and the more accurate that data, the faster the rate of genetic gain within a breeding program. Unfortunately the cost of collecting trait data from crop varieties has either remained constant or increased per data point. One way to improve the efficiency of this process is to focus on better, more precise, and faster extraction of meaningful information from the collected data.

High throughput phenotyping technology is an umbrella term which describes a wide range of new approaches that leverage advances in engineering and computer science that seek to address this current bottleneck. Generally new high throughput phenotyping technologies are approaches to measuring plant phenotypes that are predicted to be either 1) lower cost per data point, 2) more accurate, and/or 3) enable the measurement of yield relevant traits which it is not presently feasible to score.

Early infrastructure investments in high throughput phenotyping focused on automated data acquisition in controlled environment plant growth facilities (Fahlgren et al., 2015; Junker et al., 2015; Ge et al., 2016). In these facilities plants are moved around in conveyor belts (Fig. 1 a) and on regular intervals are brought into a series of imaging chambers (Fig. 1 b) where they are photographed from several angles using different types of cameras (see Fig. 1 d). A number of software tools have been developed to extract different phenotypic data from the individual 2D images generated by these automated controlled environment phenotyping facilities (Gehan et al., 2017; Lobet, 2017).

**FIGURE 1.**
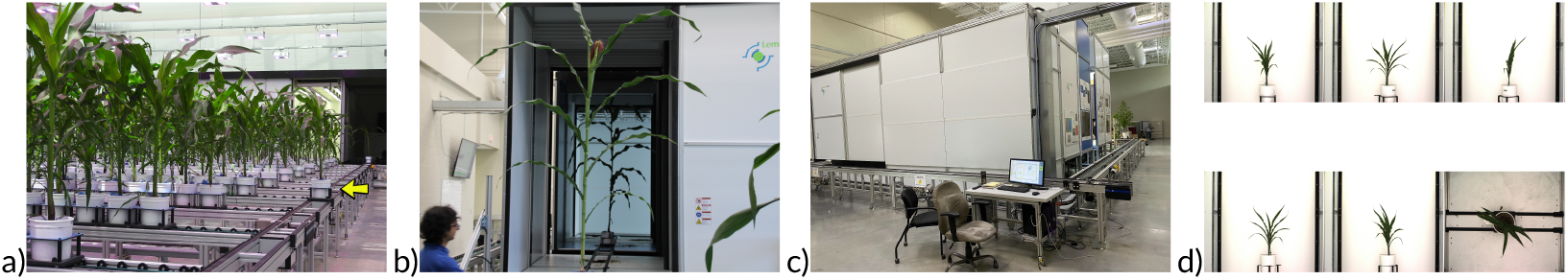
University of Nebraska Greenhouse Innovation Center phenotyping facility: a) a greenhouse that stores and automatically mobilizes plants. A plant on the conveyor belt is marked with an arrow, b) plant entering a photographic chamber, c) control unit with a computer that manages the entire facility, and d) sample image data collected from a single plant on a single day.

One key set of features linked to yield which are difficult to accurately quantify from 2D images are traits linked to canopy architecture, including leaf number, leaf angle, leaf length, internode spacing, etc. Within-species variation in radiation use efficiency, water use efficiency, and crop yield have all been linked to differences in canopy architecture (Westgate et al., 1997; Maddonni et al., 2001; Hammer et al., 2009). A large proportion of the yield gain per unit area over the past half century has come from increased planting density and breeding for lines which can thrive at higher planting densities (Duvick, 2005). In maize, selection for yield at high density indirectly selected for lines with more erect leave angles, spreading the same amount of incident light over a large photosynthetic surface (Duvick, 2005; Pendleton et al., 1968; Pepper et al., 1977). In sorghum, breeders have selected for large effect mutations with reduce internode spacing, producing a denser canopies closer to the ground (Quinby et al., 1953). Future efforts to engineer efficient canopy architectures will require quantification and simulation of a range of canopy architectures achievable from natural genetic variation in crop species (Marshall-Colon et al., 2017; Benes et al., 2020). Collecting the data needed for this objective in turn requires more comprehensive collection and characterization of genotype to genotype variation in 3D canopy architecture on a large scale.

However, plants are complex 3D structures and data collected from a single or several 2D images can miss or inaccurately estimate for important plant features (McCormick et al., 2016; Thapa et al., 2018). Various approaches attempt to faithfully extract either the full 3D structure of plants or at least some important traits from captured data. LIDAR scanning using a sensor mounted on a robotic arm has been used to reconstruct 3D models of barley plants, achieving *R*^2^ = 0.96 with ground truth measurements for predicting the area of individual leaves (Paulus et al., 2014). LIDAR-based reconstruction of maize and sorghum plants using a fixed LIDAR sensor and plants were placed on a rotating platform achieved 0.92 ≤ *R*^2^ ≤ 0.94 for maize and sorghum (Thapa et al., 2018). A time-of-flight cameras (*e.g*., Microsoft Kinect) were employed to generate 3D models of individual plants from a sorghum RIL population, achieving *R*^2^ = 0.85 with destructively measured leaf area (McCormick et al., 2016). However, these approaches require dedicated equipment unlikely to be available in many plant research labs and LIDAR are not suitable for data with high frequency information and high inter reflections that are typical for vegetation. Small parts can be easily missed and the reconstruction algorithms often fail on small or slim parts such as leaves. Conventional RGB camera are easily accessible and a number of faculties around the world use conveyor belts and rotating platforms to image individual plants from multiple angles at regular intervals (Fahlgren et al., 2015; Junker et al., 2015; Ge et al., 2016; Yang et al., 2014). One of the most commonly used method is Structure From Motion (Quan et al., 2006; Lou et al., 2014; Tomasi and Kanade, 1992) (SFM) that requires large number of images (hundreds), long processing times (minutes to hours) and often fails on texture-less or highly specular surfaces and on data that include high frequency noise, such as foliage. Although significantly more accessible than LIDAR, current approaches to 3D reconstruction from RGB images share the problems of reconstruction from LIDAR data. Other approaches to 3D reconstruction attempt to generate certain parts of plants: for example generalized cylinders (Tan et al., 2007, 2008), skeletons (Du et al., 2019), or use optimization to find a generative model with a database of leaves (Ward et al., 2015), and even reconstruct the entire growth model (St’ava et al., 2014). However, these methods usually require human interaction, perfect input data, clear background, or other requirements that make them difficult to apply in phenotyping where hundreds of plants are scanned over large periods of time.

Another well-established method is space carving of Kutulakos and Seitz (2000) that reconstruct 3D voxels occupied by the captured plant. Its main advantage is that it requires less images and a lower processing time than SFM. The drawback is that the algorithm needs an extremely precise calibration and segmentation of the object to reconstruct whereas SFM estimates calibration automatically by matching keypoints between views. Space carving is suitable to phenotyping facilities because the environment is controlled, which eases calibration and segmentation. Recent contributions focus on seedlings, which are smaller plants that are easier to reconstruct (Koenderink et al., 2009; Klodt and Cremers, 2015; Golbach et al., 2016). Other contributions focus on accelerating voxel carving with octrees (Scharr et al., 2017). Regarding 3D reconstruction, one drawback is that the algorithm does not output a surface but the photo hull of the plant, which is the maximal photo-consistent volume in which the actual plant is contained. Automatic measurements are not straightforward with a voxel representation (Golbach et al., 2016).

Calibration and segmentation are crucial to the success of space carving. For calibration, the work of Li et al. (2013) is of particular interest, because it enables calibrating of multiple cameras with non-overlapping views, which is the usual case in rotating imaging platforms. As for segmentation of plant pixels in 2D images, it can take a number of approaches. One such approach is “differencing” when the image is compared to a reference image taken by the same camera in the same imaging chamber without a plant present (Choudhury et al., 2019). A second widely adopted approach is excess green thresholding (Gehan et al., 2017). Supervised classification algorithms which consider both pixel RGB values and data from immediately adjacent pixels have been shown to exhibit higher accuracy in plant pixel segmentation than many thresholding or “differencing” based methods (Adams et al., 2020). For example, the work of Donné et al. (2016) employs a convolutional neural network to perform a segmentation.

Our contribution is the validation of a 3D reconstruction pipeline for a very low number of images: only 5 side and 1 top images. The approach automatically generates 3D voxel grids approximating the shape of the scanned plant (Fig. 2 h). This approach builds on voxel carving, but aims to achieve fully automatic reconstruction for large numbers of plants and does not require any user interaction on a per plant basis. This scalability and lack of required human interaction enabled us to evaluate the method on hundreds of plants, more than an order of magnitude greater than many previous studies. We bench marked 3D extraction and reconstruction as taking less than one minute per plant to create a 3D voxel grid with a resolution 512^3^ on a workstation equipped with an Intel Xeon W-2145 (8 cores at 3.7 GHz), with significant decreases in per plant processing time for processing multiple plants in series. We extended the voxel carving algorithm to favor recall over precision and thus get smoother plant shapes for a subsequent skeletonization. We studied the effect of the camera setup on the accuracy of the reconstruction and showed that some setups are more efficient for the same investment in money. We evaluate the impact of an imprecise calibration on the resulting 3D plants. Finally, we showed that a 3D representation brings substantial information compared to only 2D images. Our algorithm allows for quantification of traits light interception efficiency. Scoring large numbers of plants enabled quantitative genetic approaches to identifying loci controlling variation in three dimensional traits. A number of inherently three dimensional traits were estimated from 3D reconstructions from 336 sorghum plants. These trait values, combined with information on hundreds of thousands of genetic markers, were used to estimate the proportion of phenotypic variance for each trait controlled by genetic factors (heritability). For several high heritability traits it was possible to conduct genome wide association studies (GWAS) and identify specific regions of the genome controlling between plant variation.

**FIGURE 2.**
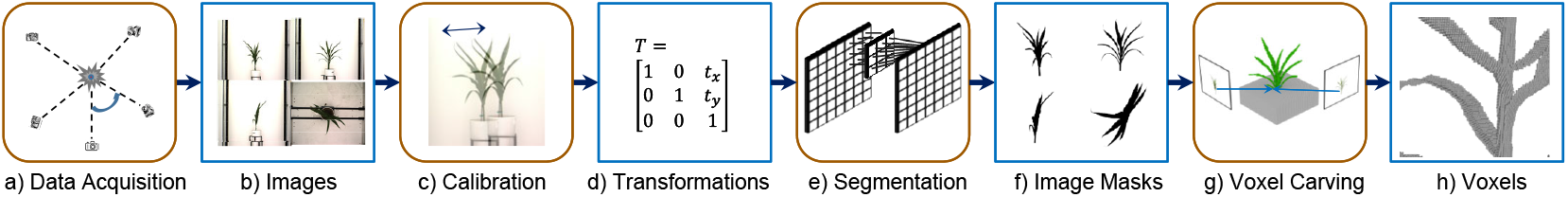
Reconstruction overview. (round boxes are processes and squared boxes are data): a) the data acquisition phenotyping chamber produces images that b) are calibrated so that the plants are centered. c)The calibrated images are segmented into binary masks by using convolutional neural network to separate the plant from the background and the masks are used to generate 3D voxel representation by d) using voxel carving.

## 2 MATERIALS AND METHOD

### 2.1 Plant Materials and Image Acquisition

Image data was acquired at the University of Nebraska Greenhouse Innovation Center’s automated phenotyping facility (Ge et al., 2016) shown in Fig. 1. Sorghum genotypes were taken from the Sorghum Association Population (Casa et al., 2008). The detailed growth conditions for the sorghum plants grown and imaged in this study were previously described in the study by Miao et al. (2020a).

Each plant was photographed and RGB images were collected from five side angles around a 360*°* with photos taken at 0*°*, 72*°*, 144*°*, 216*°*, and 288*°* plus one additional image from the top (see Fig. 1 d)). Each image had a resolution of 2,454×2,056 pixels. Plants were oriented so that the zero degree angle photo corresponded to the angle at which most leaves were perpendicular to a line between the camera and the primary stalk of the plant. The camera model is a Basler pia2400-17gc with a Pentax TV zoom lens c6z1218m3-5. Images used in this study were captured on April 11th, 2018 47 days after planting. This dataset included 2,106 distinct images and was 17.5 GB in size.

At the used distance from the camera, resolution, plant size and the zoom level, each pixel represented an area of approximately 1.56 mm^2^ for objects in the range between the camera and the pot containing the plant. Moreover, we also generated approximate 3D mesh models of corn plants using Plant Factory Exporter (v2016 R3, e-on Software) with the “maize/corn” module (Miao et al., 2019). The meshes were used for validation of the voxel carving algorithm by computing the precision and recall algorithm as discussed in Sect. 3.

### 2.2 Method

#### 2.2.1 3D Reconstruction Overview

Our method works in four steps shown in Fig. 2, where round boxes are processes and squared boxes are data. The individual steps are: 1) data acquisition, 2) calibration, 3) segmentation, and 4) voxel carving. The input are images taken from the photographic chamber of the phenotyping facility and output of our algorithm is a 3D voxel representation of the plant. In this section we provide an overview of our reconstruction pipeline that is then explained in details in following sections.

**Data acquisition** is the process of taking images of a plant (see Sect. 2). The input are the physical plants and the output are plant images. This process is fully automatic, each plant has associated unique identifier, and the images are stored on a local data server.

##### Calibration

To get an accurate reconstruction, we define a 3D coordinate system that is common to all images. A chosen reference point is the center of the pot that holds the plant. However, although the phenotyping facility attempts to align and center the pots on the rotational axis of the turn table when taking the photographs, in practice they are usually off of the main axis by up to several centimeters. This causes the pot, and thus the plant, to be misaligned in the images. Since the alignment is one of the most critical condition for the voxel carving algorithm to work well, we calibrate the images so that the pot is at the right location in every image. The results of the calibration step is a transformation matrix **T** that describes the translation that center the pot in each image.

##### Segmentation

In the next step we separate the plant from the image background. Although the lighting conditions are controlled, the varying amount of plant mass and their geometry makes the images difficult to segment by using traditional methods, such as color segmentation or thresholding. Instead, we used a convolutional neural network that is invariant to changing light and works well with soft edges that are common in plants.

##### Voxel carving

Having the images segmented from the background and knowing the pose of the cameras, we reconstruct the plant by using the voxel carving algorithm, which is a variant of the space carving algorithm of Kutulakos and Seitz (2000). The output is a 3D voxel grid that represents the photo hull of the plant, which is the maximal shape in which the true plant lays. Although it is noticeably thicker than the plant, and does not include color information, it is still well-suited for the heritability analysis (Sect. 3.4) and for other tasks such as counting the number of leaves and measuring their lengths.

##### Terminology

Let *I* denote the sequence of six input RGB images (see Fig. 1 d) *I* = [*I*_0_, *I*_72_, *I*_144_, *I*_216_, *I*_288_, *I_top_*], where the first five images [*I*_0_, *I*_72_, *I*_144_, *I*_216_, *I*_288_] are taken from the sides with angles *α* in *A* = [0*°*, 72*°*, 144*°*, 216*°*, 288*°*] (see Fig. 2 a)), and *I_top_* is the image taken from the top at an angle of 0*°*. Moreover, **P** = [**P**_0_, **P**_72_, **P**_144_, **P**_216_, **P**_288_, **P**_*top*_] refer to the corresponding projection matrices of the camera while taking images *I*. To project a 3D point *v* on the camera with angle *α*, we multiply its coordinates by the projection matrix: 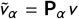, where 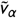 is a 2D point in image *I_α_*.

We denote the binary masks that result from the segmentation of *I* by *M* = [*M*_0_, *M*_72_, *M*_144_, *M*_216_, *M*_288_, *M_top_*] (Fig. 2f)). If the value of a pixel (*k*, *l*) in the mask *M_α_*[*k*, *l*] is equal to one, the corresponding pixel in the image 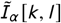 belongs to the object, while a value of zero indicates the projected position is in the background.

The final result of our algorithm is a binary voxel grid 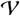 that has a resolution of 512^3^ and we refer to each voxel as *v_i,j,k_* where 0 ≤ *i*, *j*, *k* ≤ 512 denote its discrete coordinates. A voxel *v_i,j,k_* having a value one identifies a 3D position belonging to the plant photo hull, zero value identifies an empty space.

#### 2.2.2 Image Calibration

The phenotyping facility uses high precision robotic plant transportation conveyors shown in Fig. 1 a), but this precision is not sufficient to guarantee the plant to be perfectly centered in each image. The voxel carving algorithm is highly sensitive to precise location and so we perform additional image transformation to make sure the plant is centered in every image *I*. The two main sources of imprecision are 1) the pot is mounted on a turntable, but is not perfectly centered on its axis of rotation and 2) the camera’s optical center is shifted from the axis of rotation of the turntable. This causes the plant to be off-axis when rotated.

To calibrate the images, we created a 3D virtual model of the phenotyping photographic chamber by using a 3D modeling software. We then rendered a set of perfectly calibrated synthetic images of the virtual chamber without the plant, but with the pot at the exact center: one from the side and one from the top. These calibration images show how the pot should look in a theoretically perfect photographic chamber. We then used the images to find the translation that needs to be applied to the real images *I* in order to center the plant. We take a picture of the synthetic empty pot and we found its location in the calibration image by overlaying it using transparency. Then, we look at the pot in each image *I_α_* and we use phase correlation to first shift the entire picture, finally we refine the transformation with template matching. This provides the transformation matrix **T**_*α*_ that translates the image *I_α_* so that the pot is centered. The corrected image set is denoted as 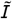 and it is calculated by transforming the input image:

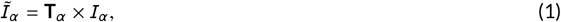

where × denotes the transformation of each individual pixel by the transformation matrix. In our dataset, we found that we had to shift images by up to 100 pixels horizontally in order to center the pot.

Our calibration only translates the images and we did not need any other optical corrections. The sorghum plants are in the size range of meters and the high-quality camera setup provides images with precision in the range of millimeters. The camera is located 5.5 meters away from the plant, which lessens perspective distortion. Moreover, the camera has a high quality optic, which limits other distortions such as vignetting.

Fig. 3 a) shows the input to the calibration step. The plant is not centered, and running voxel carving on this image would result in an imprecise reconstruction. Fig. 3 b) shows the calibration image with an empty pot that is pixel-exact centered. Fig. 3 c) shows the result of application of Eqn (1) on *I*_0_ where the input image is shifted so that the pot is at the same location as in the calibration image.

**FIGURE 3.**
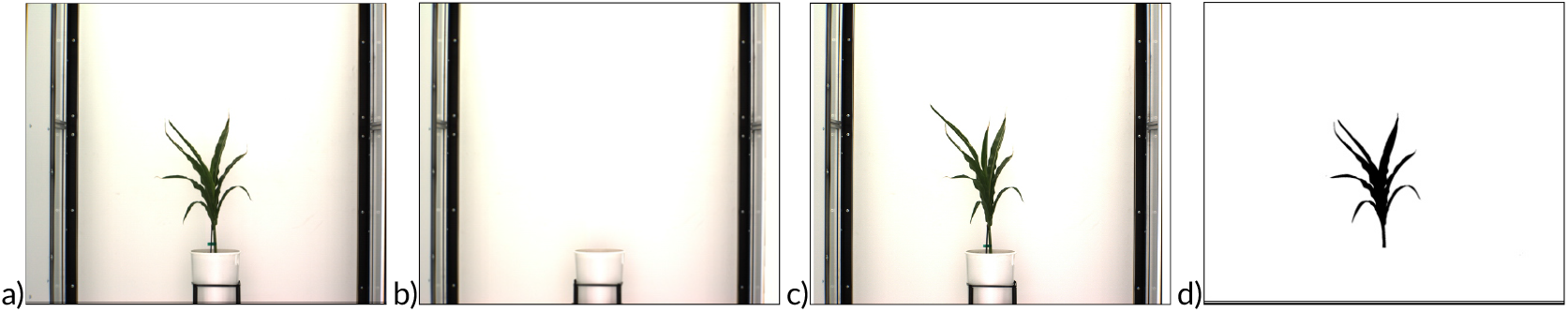
Calibration and segmentation: a) Input image *I*_0_, b) reference image from the virtual 3D chamber, c) the output corrected (centered) image 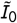, and d) the segmented mask *M*_0_.

#### 2.2.3 Image Segmentation

Although, the imaging chamber is a controlled environment that removes a lot of variability from images, they still cannot be directly input to the voxel carving algorithm because of the varying light intensity. While the lights in the chamber are fixed and constant, the plant 3D structure and complex interreflections cause huge variability in the lighting within the chamber. Therefore, it is beneficial to separate the plant in the image 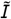 from the background.

Various techniques for image segmentation exists such as color thresholding (Lim and Lee, 1990) and color-based segmentation algorithms (Cheng et al., 2001; Haralick and Shapiro, 1985). However, they often struggle to segment the top view because they fail to differentiate between a self-shadow on the plant and the soil in the pot. Moreover, they tend to lose the leaf boundary where the color gradient changes slowly.

After experimenting with an image processing software, we noticed that good results are generated manually by varying color mapping curves and applying different color-conversion filters. However, this work is tedious, cannot be automatized, and we were not able to find a fixed set of values that would work for all plants. Eventually, we used a convolutional neural network that was trained on human-segmented images. The machine learning approach takes in account a small neighborhood around pixels when segmenting them. The training set contains 24 images, taken from four different plants with varying lighting conditions that were manually segmented. We used data augmentation to increase the variability of the dataset, in particular rotations of up to 90 degrees, shift by up to 10% of the image size, zoom in by up to 10%, and horizontal and vertical flips. While having a larger dataset would be beneficial, the manual segmentation is a tedious and lengthy process, and we found that our data set was providing good results.

The input to the neural network is a calibrated image 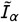 and the output is a binary image *M_α_* of the segmented plant (Fig. 3 d)). We use an architecture with four convolutional layers similar to the work of Donné et al. (2016). Our neural network (shown in Fig. S1) does not include pooling layers, which means that our images are not down sampled during processing. When classifying a pixel, our network looks at a small neighborhood around it, applies non-linear operations, and outputs a probability of belonging to the background.

Similar to the work of Milletari et al. (2016), we use the Dice loss for training because it is adapted to segmentation problems with imbalanced classes. The Dice loss is derived from the Dice similarity coefficient, which is commonly used for image segmentation validation. Having two shapes that are compared for similarity with the Dice coefficient, a value of 1 indicates perfect overlap whereas a value of 0 indicates no similarity.

We split the dataset into 18 images for training and 6 images for validation. We trained for 2,000 epochs with the Adam optimization algorithm, on a workstation equipped with a Xeon W-2145 (8 cores at 3.7GHz), 32 GB of RAM and a Nvidia TITAN Xp with 12 GB of RAM. The training took about three hours, most of which spent on data augmentation.

#### 2.2.4 Voxel Carving

The input to the voxel carving algorithm is the set of binary mask images *M* with known camera projection matrix *P*. The output is a set of voxels that correspond to the plant. Our reconstruction algorithm is voxel-based volume carving that is a variant of the volume carving algorithm of Kutulakos and Seitz (2000). Our algorithm ignores color information in pixels (texture) and focuses only on geometry.

The plant is immersed into a uniform 3D volumetric grid denoted by 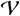 with a resolution 512^3^. The grid has a physical size 1 × 1 × 1*m*^3^ and at the resolution of 512^3^ each voxel corresponds to a volume of about 2 × 2 × 2*mm*^3^. Having a finer grid would largely increase the processing time without bringing significantly more information.

A voxel center 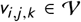 is projected on each mask image *M_α_* by using the projection matrix **P**_*α*_. A voxel is set to one *v_i,j,k_* ← 1 *i.e*., it belongs to the plant, if the projected voxel hits pixel in each *M_α_* that has also value set to one. If the voxel is projected onto at least one background pixel of the mask images (value zero), it is set to background *i.e*., *v_i,j,k_* ← 0:

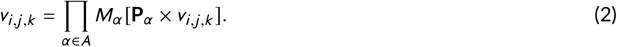

Fig. 4 shows schematically this process for two masks rotated by 90*°* and denoted by *M*_0_ and *M*_90_. The voxel *v_i,j,k_* is projected to *M*_0_ by using the projection matrix **P**_0_ and to *M*_90_ by using **P**_90_. If the corresponding pixels in both masks are equal to one, the voxel *v_i,j,k_* is set to one.

**FIGURE 4.**
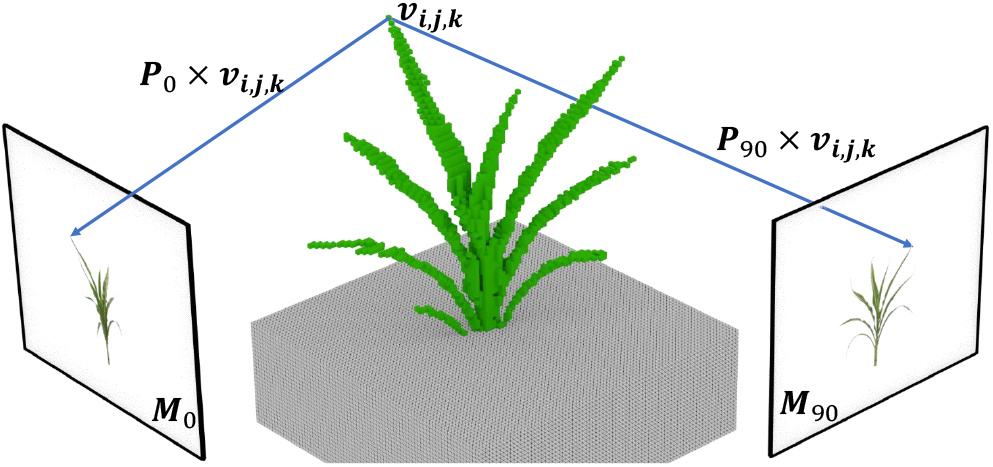
Illustration of the voxel carving algorithm employed in this study. The voxel carving algorithm projects each voxel *v_i,j,k_* to the corresponding masks (*M*_0_ and *M*_90_ in this example) by multiplying its center by the calibrated projection matrix **P** indicated as rays. If the projected voxel’s position in all masks corresponds to pixels with value one the voxel is also set as to one *i.e*., as being a part of the plant.

The algorithm has a complexity of at least 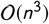 with *n* being the number of the voxels of the grid. However, this algorithm is also embarrassingly parallel and it can process as many voxels in parallel as available processing units. Its output is a voxel grid 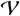 that associates to each voxel *v_i,j,k_* a binary value indicating whether it is a part of the plant photo hull or not (the photo hull is the maximal shape that the plant occupies).

Voxel carving is highly sensitive to camera calibration. In fact, if one part of the plant is missing or distorted in only one image, it will not be included in the reconstructed volume (Eqn (2). This causes problems with thin parts, such as the leaf tips or leaves projected from side.

To address this issue, we extended the voxel carving algorithm by looking at a small neighborhood in images instead of only a single pixel. When processing a voxel, we project a sphere of a fixed radius *R* on each input binary image *M_α_*. If the projected sphere covers some pixels in the mask belonging to the plant, it is considered a match:

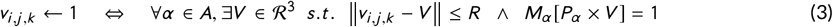

This improves the recall of the reconstruction at the expense of precision as shown in the validation in Sect. 3.3. However, it is better to have higher recall values, because it corresponds to plant with continuous areas that are easier to post process. When using the sphere instead of a single value, the reconstructed plant is more faithful and includes less holes. The reconstructed leaves are thicker, but it better captures the overall plant shape. In other words, leaves are still at the correct location and their length and width are captured correctly as shown in Fig. 5. Moreover, in order to make the reconstructed plant connected, we remove voxels that do not belong to the major connected component *i.e*., voxels hanging in space.

**FIGURE 5.**
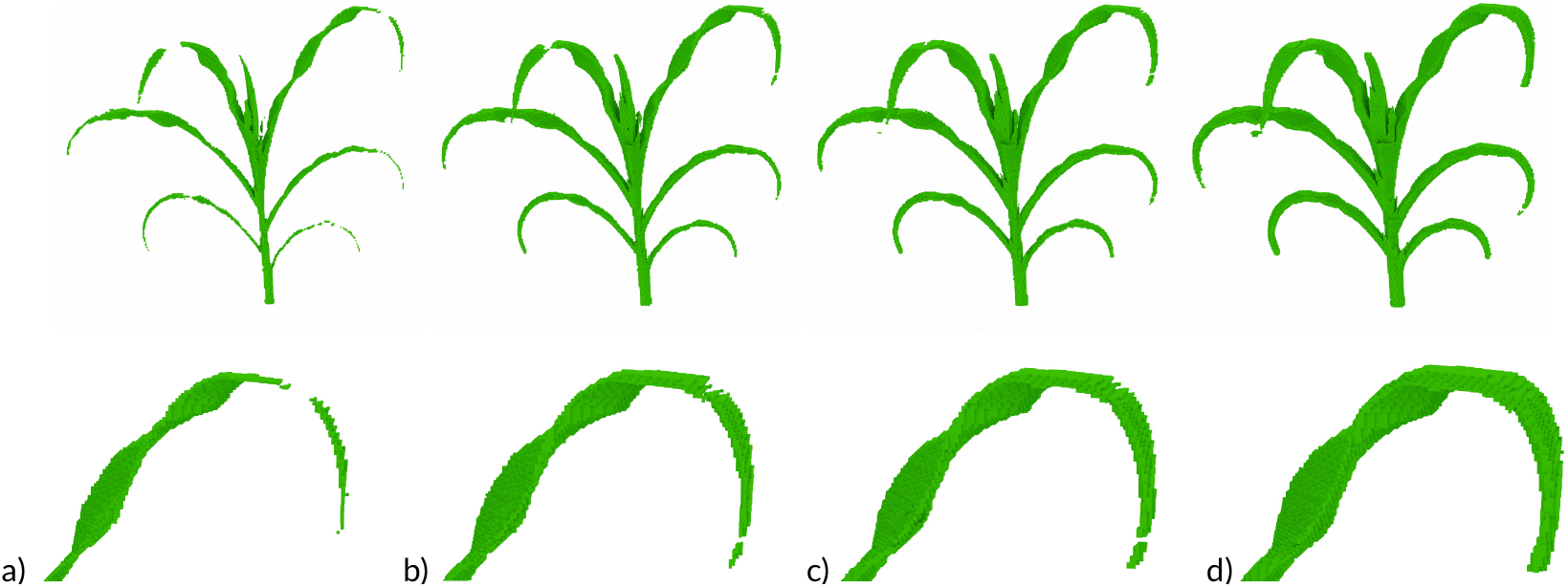
Effect of varying the size of the sensitivity area on the final 3D reconstruction of plant structures. Reconstructions of either an entire plant (top) or detailed view of the reconstruction of the upper right leaf from the same plant (bottom): when the sensitivity area is set to a sphere with a radius a) one, b) two, c) three, and d) four centimeters.

#### 2.2.5 Trait Extraction and Genome-Wide Association Studies (GWAS)

The reconstructed voxels were used to quantification of four traits for each sorghum line in SAP. The number of voxels in a reconstructed plant which approximates the plant volume is denoted by

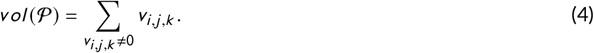

where 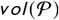 is the volume of a plant 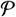 represented by the total number of voxels. The volume of the bounding cylinder was also calculated as an approximation of the space occupied by a plant. Each plant was encapsulated into its bounding cylinder which approximates the space occupied by a plant. The volume of bounding cylinder is calculated in cylindrical coordinate system and it is tighter compared to the bounding box system. We constrain the axis of the bounding cylinder to be coincident with the *z*-axis.

Moreover, we also calculate the shadow area caused by light arriving from the top of the plant and it was denoted by 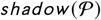 and calculated as

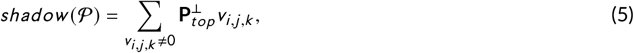

where 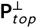 is the top orthogonal projection matrix. If a projected voxel falls into shadow it is counted only once. The value of 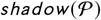 is directly proportional to the cosine-corrected amount of intercepted light arriving from the top which approximates the amount of sun energy received by a plant (Benes, 1997; Soler et al., 2003).

The ratio of the number of voxels to the shadow area indicates the energy use efficiency 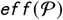 by a plant 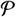 and it is calculated as

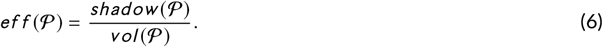

Trait values described above where combined with a published set of 569,306 SNP markers for the sorghum association population (Miao et al., 2020b) using the mixed linear model (MLM) based GWAS algorithm implemented in GEMMA using mixed linear model (MLM) (Zhou and Stephens, 2012). The first three principal components calculated from Tassel of Bradbury et al. (2007) were fit fixed effects. A kinship matrix calculated within GEMMA was fit as random effect. The number of independent SNPs was estimated using the GEC/0.2 software package described by Li et al. (2012). A Bonferroni corrected p-value of 0.05 based this estimated number of independent SNPs was used as the cut off for statistically significant SNP-trait associations. The phenotypic variation explained (PVE) value reported by GEMMA was used to estimate the narrow-sense heritability for each trait.

## 3 RESULTS AND DISCUSSION

### 3.1 Segmentation Validation

The average Dice similarity coefficient between the predicted validation set and the ground truth images was about 0.997 in our experiments. We visually inspected the masks and the segmentation in all images and the result is correct except when the plant has a support that is sometimes classified as background even if it is over the plant, which cuts the leaf in the reconstructed plant. We also noticed in some top views that some dirt stains on the floor may be misclassified as belonging to the plant. However, having wrong pixels only in one view is rejected in the voxel carving algorithm that uses voting mechanism (Eqn (3)). An artifact that would be classified as a voxel would need to be in all views simultaneously that is virtually impossible. Although a deeper neural network could potentially provide better results, we faced a memory limitation while training on our GPU. The images are large, which causes two problems: 1) the dataset is larger that the available RAM and 2) we could not add many layers and filters. A possible avenue for future work is to training on the CPU with more RAM. Moreover, we could use the Tversky loss (Salehi et al., 2017), which is a generalization of the Dice loss, to favor recall over precision. In fact, when it comes to voxel carving adding superfluous information in images is less problematic than removing essential information.

### 3.2 3D Reconstruction Validation

The accuracy of plant 3D reconstruction was assessed using synthetic data generated from procedural maize plants and an *in silico* reconstruction of the plant imaging chamber at the University of Nebraska-Lincoln by using the dataset of Miao et al. (2019).

Ten triangle meshes of maize plants has been generated using Plant Factory Exporter (see examples in Fig. S2). We visually inspected the 3D models to makes sure that they did not include self-intersections or other errors. The generated plants had 11 leaves on average, the minimum and maximum were 9 and 15 respectively. The ten meshes were voxelized and the voxel grids were used as ground truth for 3D reconstruction.

We then simulated the data acquisition step by rendering six images per plant, five side and one top view, by replicating the parameters employed for real-world data collection. These images were already calibrated and segmented since we tuned the simulation of the data acquisition to output directly the plant masks. Finally, we reconstructed the photo hull of the plants based on the 6 images and the 6 camera matrices by using our voxel carving algorithm.

We compared the ground truth voxelized meshes with the ones reconstructed by using our algorithm and we use information retrieval evaluation measures: precision, recall, and F-measure. Precision is the fraction of reconstructed voxels that are actually part of the plant. Let’s denote by *t p* (true positive) the number of detected voxels that are present in both grids - the ground truth and the reconstructed one. Let’s denote by *f p* (false positive) the number of detected voxels present in the reconstructed grid but not in the ground truth, and let’s denote *f n* (false negative) the number of detected voxels that are present in the ground truth, but were not reconstructed. We define precision *p* as

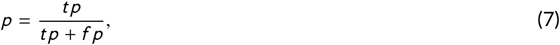

recall is denoted by *r* and it is the fraction of voxels from the actual plant that were successfully reconstructed

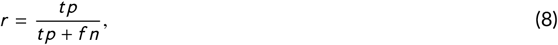

and F-measure *F m* combines precision and recall and it is the harmonic mean of both measures

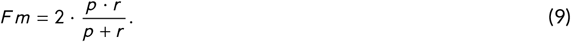

We calculated *p*, *r*, and *F m* for the ten maize models and we also calculated the average and standard deviation. This gives an upper bound on the accuracy we can get using voxel carving using our particular camera setup and the results are in Table 1.

**TABLE 1.** Precision, recall, and F-measure of the 10 synthetic plant reconstructed by using our algorithm.

| measure | average | standard deviation |
| --- | --- | --- |
| precision $p$ | 0.4985 | 0.0429 |
| recall $r$ | 0.9530 | 0.0059 |
| F-measure $Fm$ | 0.6536 | 0.0372 |

While precision *p* ≈ 0.5, recall *r* > 0.95, which indicates that no essential parts of the plant are missing. As explained in Sec. 2.2.4, although the precision is not high, imprecision does not affect the overall shape of the plant and does not have a big impact on measurements such as plant height and leaf lengths. Fig. 6 shows visual comparison of generated plants and their reconstruction. Sec. 3.5 discusses further details on how the camera setup affects the reconstruction accuracy.

**FIGURE 6.**
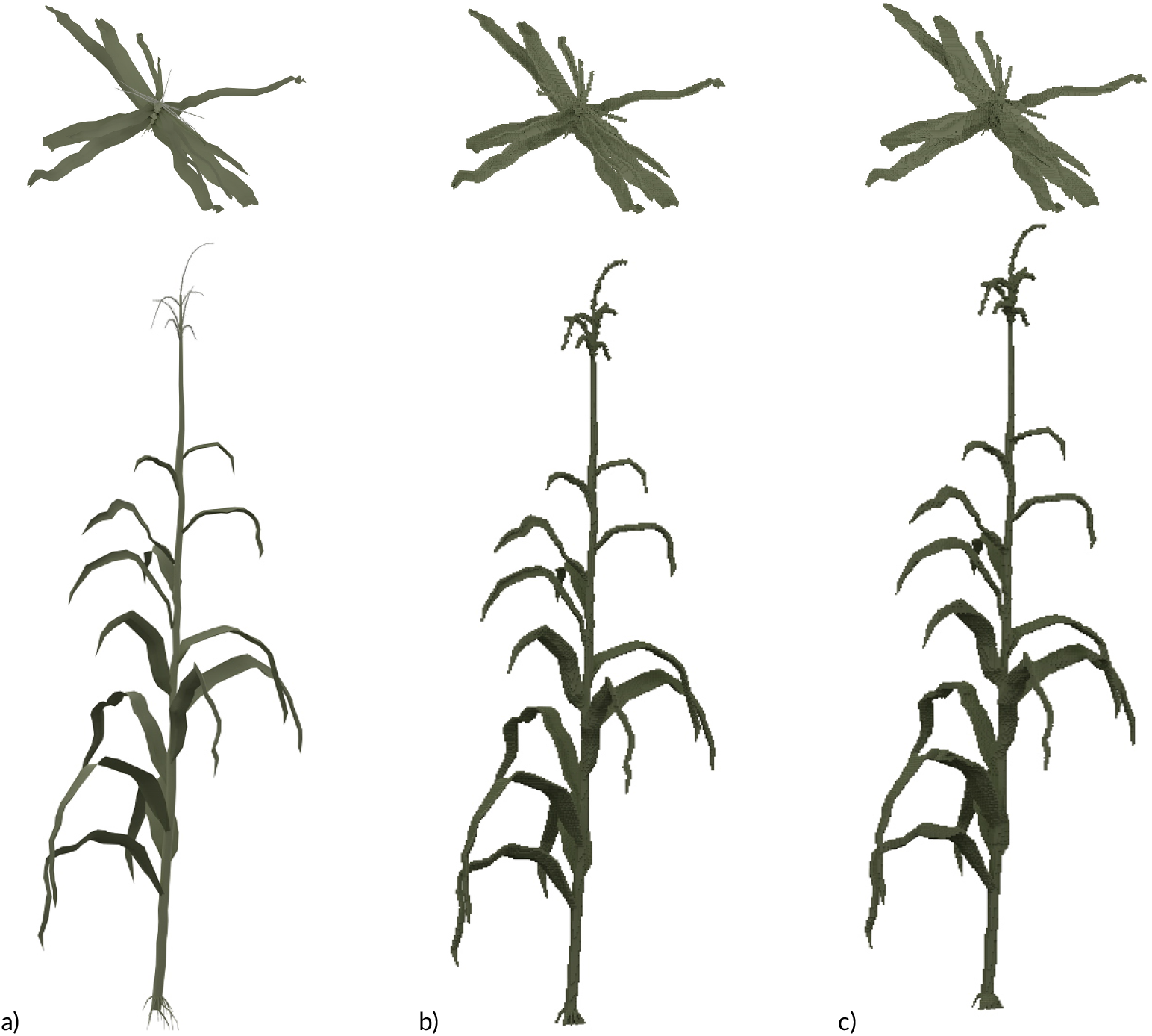
Estimating upper bound reconstruction accuracy using simulated data. Side and top views of a) a procedurally generated 3D model of a maize plant, b) a voxelized version of the original 3D model, and c) a reconstructed 3D model generated using the voxel carving algorithm describes in this paper, and six simulated 2D images generated from the original 3D model (5 side views and one top view).

### 3.3 3D Reconstruction Accuracy

We ran the reconstruction pipeline on hundreds of plants and it would be difficult to verify the accuracy of each plant manually. Inspired by the work of Klodt and Cremers (2015), we computed the Dice coefficient denoted by *D* and used it to verify whether a plant was successfully reconstructed without visually inspecting it. We re-project all voxels from each plant to the six views and we then compute *D* between the re-projected images and the plant masks obtained after the segmentation. The value of *D* = 1 indicates that everything in the plant masks has been reconstructed and *D* = 0 indicates that the reconstruction failed (see Sec. 2.2.3 for details about the Dice coefficient). In our experiments, we found an average value of *mean*(*D*) = 0.884 with a standard deviation of *stdev*(*D*) = 0.011 for the 339 non-empty plants from our data set.

To provide a visual insight why some plants have a low Dice coefficient value, we color-coded the reprojections from each camera and alpha-blended them with the corresponding RGB plant images. The results in Fig. S3 show in green true positives, blue color encodes false positives, red value depicts false negative *i.e*., a part of the plant that has not been reconstructed, and white color is a true negative.

We identified two factors that negatively affect the Dice coefficient. First, when taking pictures, leaf tips move as the plant rotates on the platform and sometimes they are not perfectly stable. Second, the segmentation step leaves small artifacts, especially on the top view, which negatively impact the Dice coefficient even if the reconstruction is accurate. Nevertheless, we can quickly identify a badly reconstructed plant when its value is significantly off the distribution (*D* < 0.8 in our experiments).

We found that common errors (see examples in Fig. S3) causing low value of Dice coefficient include a) plants with incorrectly segmented dry leaves, b) broken stems that make parts of the plant invisible from the top view or out of the voxel grid, c) failed calibration due to leaves laying in the pot in one view, d) small plants that cause noisy reconstruction, and e) dead plants (empty pots).

### 3.4 Variation in 3D Structure Among the Sorghum Association Panel (SAP)

We captured a data set for 229 Sorghum lines in sorghum association panel (SAP) at early growth stage using a high throughput phenotyping facility (Ge et al., 2016; Miao et al., 2019) (Sect. 2.2). The dataset includes a total of 1,374 (229 lines × 6 viewing angles) RGB images and has overall size of 12.0 GB. We ran the entire 3D construction pipeline on them.

The shadow area 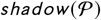 Eqn(5) and the number of voxels 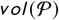 Eqn(4) follow a strong linear pattern with the correlation coefficient of 0.81 (P-value=2.8e-81) (see Fig 7 b)). Overall, sorghum plants with large number of detected voxels tend to have a higher shadow area. However, some lines out of this rule are also observed. For example, the sorghum line PI534105 has a large shadow area but a relative small number of voxels. In contrast, PI533821 has a large number of voxels but a relative small shadow area. After checking the 3D models of these two plants [Supplementary file S1], we found that the differences are results of the leaf architecture in these two plants: PI533821 has a more compact leaf architecture than PI534105 (see Fig 7 b)). Compared to the total sun energy a plant receives, looking at the energy use efficiency, *i.e*., the ratio of shadow area to the number of voxels 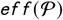 Eqn(6) is more meaningful and realistic when evaluating a sorghum line in breeding programs. As shown in Fig. 7 d, large plants tend to be less energy use efficiency compared to the relative smaller plants but there is no clear linear relationship between these two features.

**FIGURE 7.**
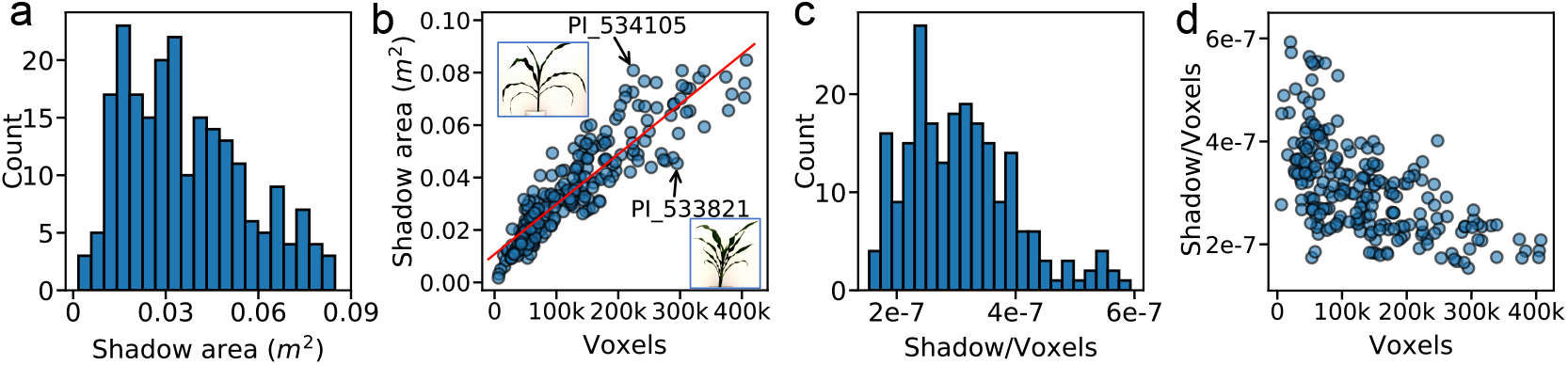
Distribution and relationship of radiation use efficiency related traits in the Sorghum Association Panel. a) The distribution of shadow area 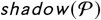 Eqn(5) across sorghum lines in the sorghum association panel (SAP) tested in this study. b) Relationship between shadow area and the number of voxels. The best-fit linear regression line is indicated in red with the equation y = 1.91e-7x + 0.01. Two sorghum lines with particularly high or low ratios are indicated with arrows and their silhouettes are shown. c) The distribution of ratio of shadow area to number of voxels across sorghum lines in the sorghum association panel (SAP) tested in this study. d) Relationship between the shadow area/voxels ratio (e.g. 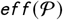 Eqn(6) and the total number of voxels for a given plant.

Narrow-sense heritability was estimated for features extracted from 3D sorghum models. The estimated narrow-sense heritability for the number of voxels, bounding cylinder volume, shadow area, and the ratio (shadow area/number of voxels) were 0.51, 0.49, 0.59, and 0.65 respectively. This results suggest that all the features mentioned above are sufficiently heritable for the mapping of individual loci controlling between genotype variation. Genome wide associations studies were conducted for each of the four phenotypes. Statistically significant trait associated SNPs were identified for both bounding cylinder volume and the ratio of shadow area to number of voxels 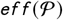 [Figure 8]. Two significant signals were identified for bounding cylinder volume [Figure 8a]. The significant peak on chromosome 7 is associated with *dwarf3*, a classical sorghum mutant which encodes a MDR transporter influencing cell elongation potentially via polar auxin transport (Multani et al., 2003). The second signal for bounding cylinder volume is a single SNP located within the gene Sobic.005G070200 on sorghum chromosome 5. Sobic.005G070200 encodes a wall-associated receptor kinase galacturonan-binding protein. Two well supported and statistically significant clusters of trait associated SNPs were identified for (shadow area/number of voxels) 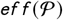 (Fig. 8 b)). One of these peaks, on sorghum chromosome 6, corresponds to a second classical sorghum dwarf gene, *dwarf2* (Hilley et al., 2017). However, the second well supported peak, on sorghum chromosome 3, is novel. To the best of our knowledge none of the genes associated with this shadow area/number of voxels 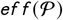 have been previously linked to functions in determining canopy architecture or leaf morphology [Supplementary file S2].

**FIGURE 8.**
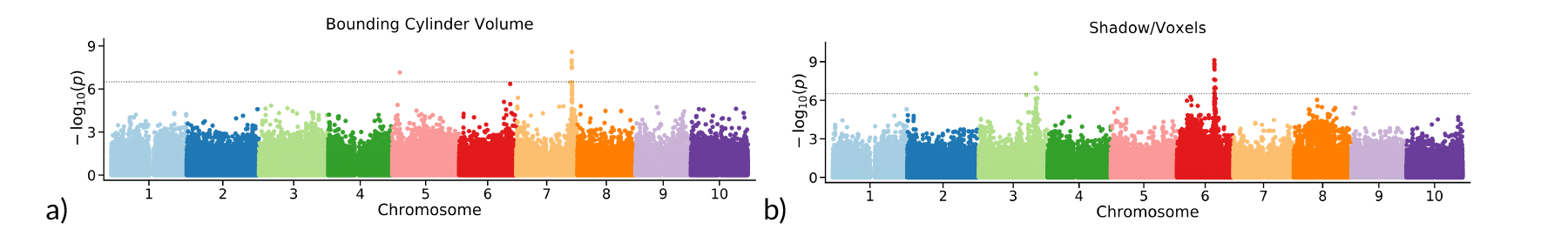
Genome wide association analyses to identify genetic loci controlling variation in 3D phenotypes. a) Manhattan plot summarizing the results of a genome wide association study conducted using bounding cylinder volumes calculated from 3D reconstructions of sorghum lines from the SAP. b) Manhattan plot summarizing the results of a genome wide association study conducted using shadow area/voxels ratio (e.g. 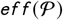 Eqn(6 calculated from 3D reconstructions of sorghum lines from the SAP. Horizontal dashed line indicates a Bonferroni multiple testing-corrected threshold for statistical significance equivalent to p-value = 0.05.

### 3.5 Optimizing Image Acquisition for 3D Reconstruction

Having a complete 3D model of the phenotyping chamber, we were able to perform a set of virtual experiments that measure behavior of the reconstruction in varying conditions and result in several suggestions for image acquisition. We use the benchmark from Sect. 3.2 to evaluate different configurations of the camera setup.

We tested three parameters and checked how they affect the 3D reconstruction: 1) the number of captured images, 2) the presence of the top camera, and 3) the presence of an optional third camera looking at the plant from the middle of the angular distance between the top and the side cameras *i.e*., at 45°. We call the configurations as *side* (side cameras), *top*, and *angle*. We compare the reconstruction accuracy when taking: 3, 5, 7, 9, 11, 13, and 15 *side* images, with or without the *angle* and *top* cameras. In total, we compared 28 different camera setups and results are shown in Fig. 9 and the values in Table S1 in Supporting Material.

**FIGURE 9.**
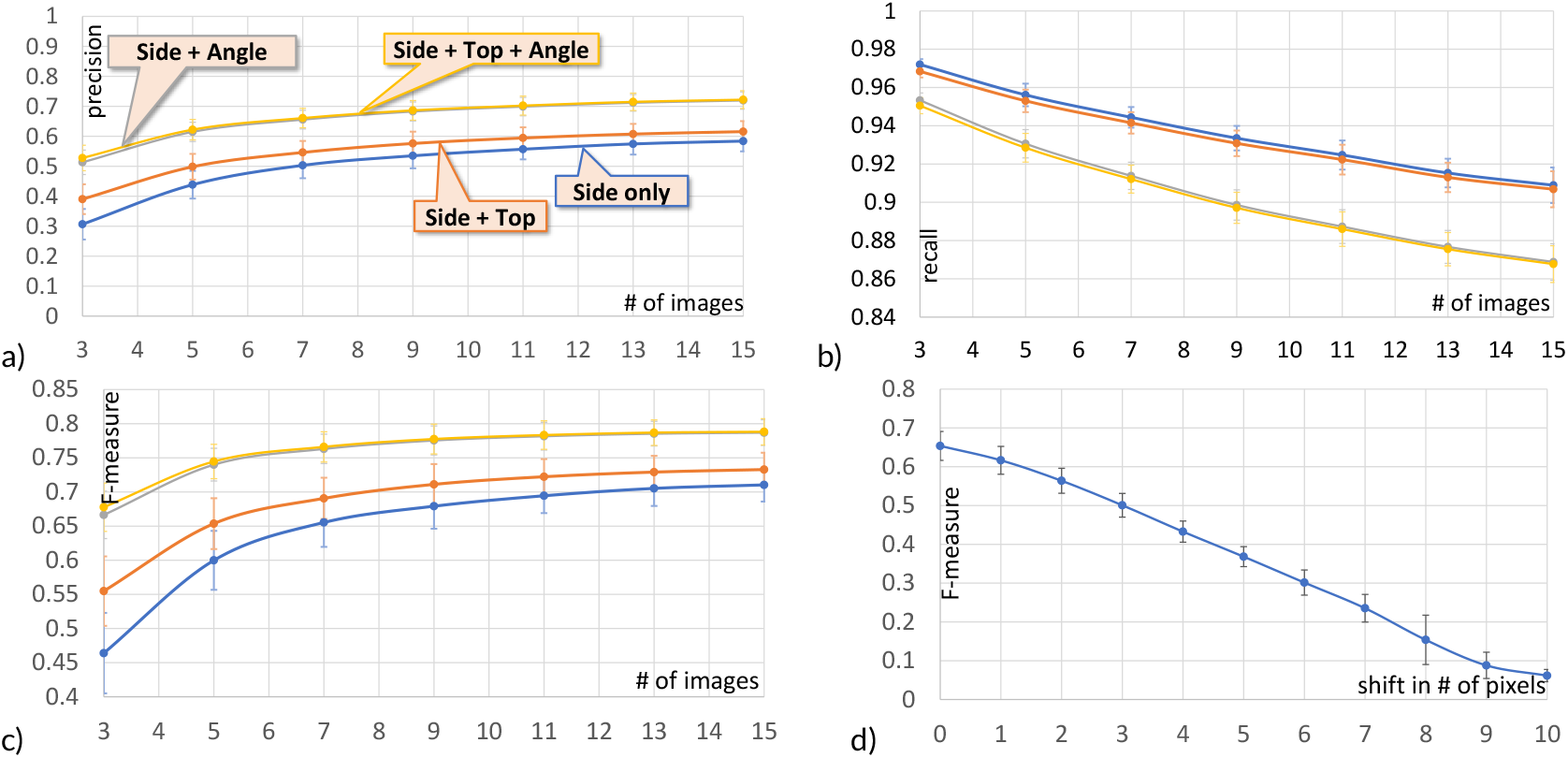
Impact of camera setup on reconstruction accuracy: a) precision, b) recall, and c) F-measure with respect to the camera setup and the number of images taken around the plant. d) F-measure with respect to the side image camera offset in pixels.

With the increasing number of captured images the precision increases at the expense of recall independently of the presence of the *top* and the *side* camera. This behavior is expected: the number of detected voxels in voxel carving can only decrease when adding a new image. In other words, we reconstruct less volume of the plant, but what we get is more precise. The F-measure always increases as we add images. Taking more images improves the reconstruction accuracy. However, the marginal benefit - quantified by F-measure - for each additional side view image plateaus at relatively low values. Adding additional cameras which observe the plant from additional angles both provides large initial increases in precision and F-measure but also reach higher values before plateauing. Adding additional cameras to provide more viewing angles increases the cost of an imaging setup but not the time for data acquisition. In contrast, collecting more side views from the same camera does not increase the cost of construction, but significantly increases time required per plant and decreases throughput, because the system needs to wait for the plant to stabilize its motion after each rotation.

The *angle* and the *top* cameras always improve the reconstruction. Fig. 9 c) shows that with a fixed number of images adding a camera always yields to a better F-measure. An important observation is that the effect on the F-measure is greater when adding the *angle* camera than the *top* camera. For the same investment (two cameras) we can get better data by setting up the second camera as *angle* instead of *top* as is the common practice. Our phenotyping facility is a closed system that does not allow us to make modifications of the setup. However, Scharr et al. (2017) used a three camera setup with a *angle* camera and they report success in plant reconstruction.

The calibration step is essential for voxel carving, because the plant is not pixel-exact centered and the *side* camera always produced shifted images. We thus ran a study to estimate the loss in accuracy due to bad calibration. In particular, we run our reconstruction while simulating an imperfect calibration by translating simulated side images along the *x*—axis by 1 to 10 pixels. We report the F-measure of the reconstruction in Fig. 9 d) and in Tab. S2 in Supporting Material. Even a shift of 2 pixels caused a drop of 14% of the F-measure (from 0.6536 to 0.5638). Shifting the image by 10 pixels caused a loss of 91% (from 0.6536 to 0.0617).

### 3.6 Based Practices When Capturing Images for Future 3D Reconstruction

Experiments in phenotyping facilities are expensive and time consuming and generally cannot be easily repeated to change or improve image acquisition. In this section, based on our experience, we provide several guidelines to best use phenotyping facilities in order to maximize the amount of information captured and improve the potential for future reuse.

When scanning plants with long leaves a good option is either the camera rotating or use multiple cameras as opposed to rotating the plant (Kumar et al., 2014). We are not aware of a reconstruction algorithm that is robust enough to deal with leaves that move between pictures. It is better to put the camera farther from the plant and change the focal length so that the plant occupies most of the image, rather than placing the camera close to the plant. For a plant occupying an equal area in an image, a distant camera with a long focal length flattens perspective distortion, reduces blur and maximizes the per-pixel precision of images relative to a close camera with a short focal length. In Sec. 3.5, we simulated and evaluated a range of imaging set ups and camera position options. Adding more images improves the result, but only up to a certain point due to a plateau effect. An alternate way to improve the reconstruction accuracy is to add more camera angles. When only one camera is available, we recommend taking side views with it. If a second camera is available, our virtual experiments show that a high angle shots provides more information than a camera looking directly down on the plant..

Colors in RGB images have a considerable importance in allowing or hindering segmentation of plant pixels from background pixels. A white background makes it easy to separate a green plant. When using supports for the plants, such as pots or a duct tape, it is best to avoid green or dark colors because are much more likely to be misclassified than bright pixels such as blue, red, or yellow. The main reason is that when using a segmentation method based on colors, it is hard to distinguish green and dark colors from the plant, as shaded parts of the plant can be closer to black than green.

When using voxel carving with only a few images - a common scenario in most phenotyping facilities - the voxel reconstruction will often contain artifacts that look like leaves. Many of these can be quickly discarded by selecting the major connected component of voxels, as many artifacts will not be connected to the real plant. However, we still observed a number of also artifacts attached to plants. These prove more difficult to remove, and future work is needed to address them. In the short term, we can only urge researchers collecting data using automated phenotyping facilities to capture as many views from as many distinct viewing angles as the logistics of their experimental design allows.

## Data Availability

The authors commit to deposit both the raw images analyzed as part of this study, voxel-based 3D reconstructions, as well as trait values for four 3D phenotypes quantified for each plant into DataDryad when and if the manuscript is accepted for publication. Genetic marker data employed in this study was previously published in Miao et al. (2020a) and has been deposited in FigShare doi: 10.6084/m9.figshare.11462469.v4

## Acknowledgements

We would like to thank Melba Crawford and Behrokh Nazeri for discussion and initial help with the data retrieval. Valerian Méline for discussion about plant biology and limits of current phenotyping and Lydia Lindner for helping successfully training the segmentation neural network. This project was completed utilizing the Holland Computing Center of the University of Nebraska, which receives support from the Nebraska Research Initiative.

## Supporting Information

Please note that we include 3D models and sample images in support material together with this paper. The 3D data are in OBJ format that can be viewed for example by using free MeshLab software (http://www.meshlab.net/)

**FIGURE S1.**
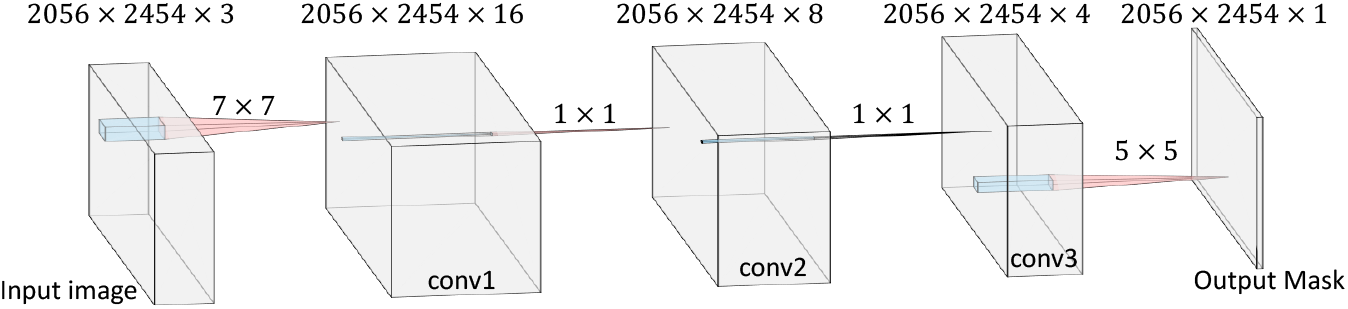
Deep neural network used for image segmentation.

**FIGURE S2.**
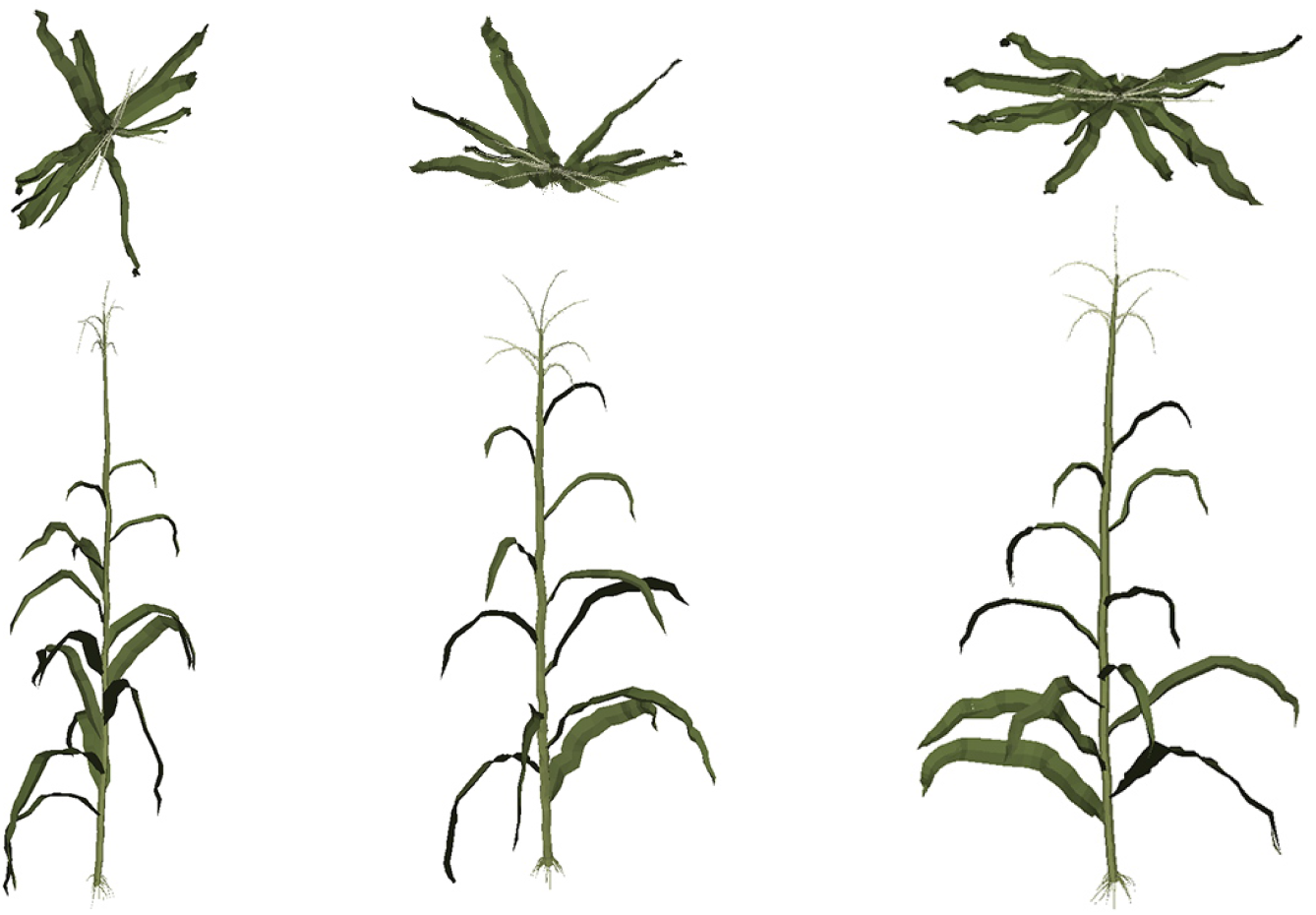
Examples of procedurally generated 3D models of maize (top and side views) used as ground truth in assessing 3D reconstruction accuracy.

**FIGURE S3.**
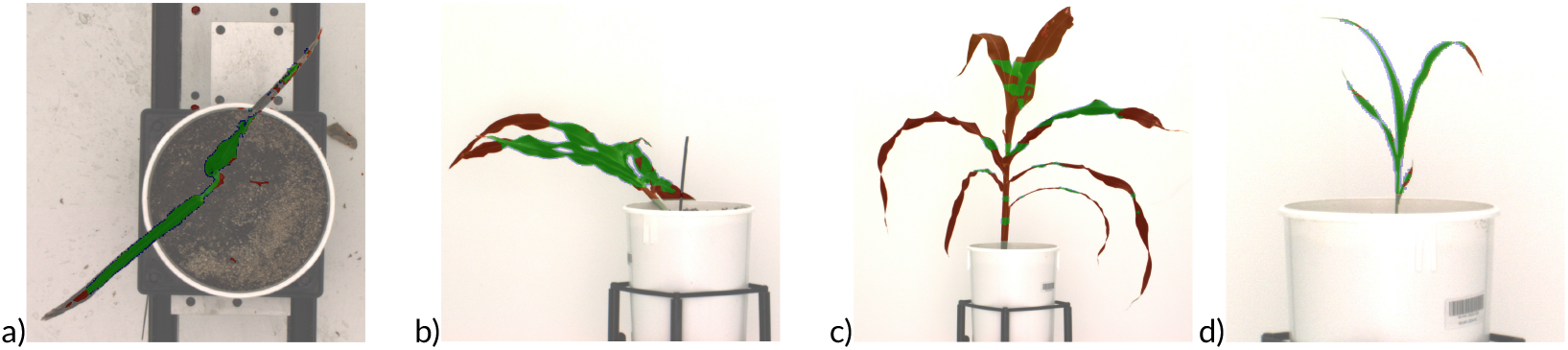
Examples of plants that were automatically detected as errors by having low value of Dice coefficient.

**TABLE S1.** Precision, recall, and F-measure of the bench marking of the virtual chamber with *top* and *angle* cameras.

| angle/top | cameras | *p* | *stdev(p)* | *r* | *stdev(r)* | *Fm* | *stdev(Fm)* |
| --- | --- | --- | --- | --- | --- | --- | --- |
| x/x | 3 | 0.3066 | 0.0508 | 0.972 | 0.0028 | 0.464 | 0.0589 |
| x/x | 5 | 0.4385 | 0.0462 | 0.9561 | 0.0059 | 0.5999 | 0.0432 |
| x/x | 7 | 0.5031 | 0.0429 | 0.9444 | 0.0054 | 0.6554 | 0.0358 |
| x/x | 9 | 0.535 | 0.0417 | 0.9335 | 0.0064 | 0.6792 | 0.033 |
| x/x | 11 | 0.5569 | 0.0338 | 0.9247 | 0.0076 | 0.6945 | 0.0254 |
| x/x | 13 | 0.5745 | 0.0353 | 0.9153 | 0.0075 | 0.7053 | 0.0258 |
| x/x | 15 | 0.5839 | 0.034 | 0.9089 | 0.0093 | 0.7104 | 0.0245 |
| x/✓ | 3 | 0.3904 | 0.0497 | 0.9684 | 0.0032 | 0.5548 | 0.0509 |
| x/✓ | 5 | 0.4985 | 0.0429 | 0.953 | 0.0059 | 0.6536 | 0.0372 |
| x/✓ | 7 | 0.5462 | 0.0382 | 0.9415 | 0.0056 | 0.6906 | 0.0302 |
| x/✓ | 9 | 0.5762 | 0.0394 | 0.9308 | 0.0067 | 0.7111 | 0.0297 |
| x/✓ | 11 | 0.5945 | 0.0355 | 0.9223 | 0.0078 | 0.7223 | 0.0259 |
| x/✓ | 13 | 0.6076 | 0.0338 | 0.913 | 0.0077 | 0.7291 | 0.0239 |
| x/✓ | 15 | 0.6157 | 0.035 | 0.9068 | 0.0094 | 0.7328 | 0.0247 |
| ✓/x | 3 | 0.5133 | 0.0408 | 0.9533 | 0.0038 | 0.6665 | 0.0347 |
| ✓/x | 5 | 0.6148 | 0.0322 | 0.9306 | 0.0074 | 0.74 | 0.024 |
| ✓/x | 7 | 0.6557 | 0.0311 | 0.9138 | 0.0071 | 0.7631 | 0.0215 |
| ✓/x | 9 | 0.6827 | 0.0316 | 0.8986 | 0.0079 | 0.7755 | 0.0212 |
| ✓/x | 11 | 0.699 | 0.0306 | 0.8873 | 0.0088 | 0.7817 | 0.0203 |
| ✓/x | 13 | 0.7118 | 0.0277 | 0.8767 | 0.0086 | 0.7854 | 0.0181 |
| ✓/x | 15 | 0.7195 | 0.0288 | 0.8688 | 0.0095 | 0.7869 | 0.0188 |
| ✓/✓ | 3 | 0.5276 | 0.0426 | 0.9505 | 0.0042 | 0.6777 | 0.0354 |
| ✓/✓ | 5 | 0.6222 | 0.0342 | 0.9285 | 0.0075 | 0.7446 | 0.0252 |
| ✓/✓ | 7 | 0.6609 | 0.0324 | 0.9121 | 0.0073 | 0.766 | 0.0221 |
| ✓/✓ | 9 | 0.6866 | 0.0328 | 0.8971 | 0.0081 | 0.7775 | 0.0218 |
| ✓/✓ | 11 | 0.7025 | 0.0318 | 0.886 | 0.009 | 0.7833 | 0.0209 |
| ✓/✓ | 13 | 0.715 | 0.0289 | 0.8755 | 0.0087 | 0.7869 | 0.0187 |
| ✓/✓ | 15 | 0.7223 | 0.0298 | 0.8677 | 0.0096 | 0.7881 | 0.0193 |

**TABLE S2.** The effect of shift of the camera on the reconstruction precision.

| Pixel shift by | 0 | 1 | 2 | 3 | 4 | 5 | 6 | 7 | 8 | 9 | 10 |
| --- | --- | --- | --- | --- | --- | --- | --- | --- | --- | --- | --- |
| F-measure | 0.6536 | 0.6165 | 0.5638 | 0.5007 | 0.4327 | 0.3682 | 0.3014 | 0.2354 | 0.1538 | 0.0881 | 0.0617 |

